# Direct observation of Hsp90-induced compaction in a protein chain

**DOI:** 10.1101/2021.08.08.455546

**Authors:** Alireza Mashaghi, Fatemeh Moayed, Eline J. Koers, Günter Kramer, Matthias P. Mayer, Sander J. Tans

**Affiliations:** AMOLF, Science Park 104, 1098 XG Amsterdam, The Netherlands; Leiden Academic Centre for Drug Research, Leiden University, Einsteinweg 55, 2333 CC Leiden, The Netherlands; Center for Molecular Biology of Heidelberg University (ZMBH), DKFZ-ZMBH Alliance, Im Neuenheimer Feld 282, D-69120 Heidelberg, Germany; German Cancer Research Center (DKFZ), Im Neuenheimer Feld 282, D-69120 Heidelberg, Germany; Department of Bionanoscience, Kavli Institute of Nanoscience, Delft University of Technology, Delft, The Netherlands

**Keywords:** Hsp90, chaperone, protein chain compaction, conformational heterogeneity, optical tweezers

## Abstract

The chaperone Hsp90 is well known to undergo important conformational changes, which depend on nucleotide, co-chaperones, substrate interactions and post-translational modifications. Conversely, how the conformations of its unstable and disordered substrates are affected by Hsp90 is difficult to address experimentally, yet central to its function. Here, using optical tweezers and luciferase and glucocorticoid receptor substrates, we find that Hsp90 promotes local contractions in unfolded chains that drive their global compaction down to dimensions of folded states. This compaction has a gradual nature while showing small steps, is stimulated by ATP, and performs mechanical work against counteracting forces that expand the chain dimensions. The Hsp90 interactions suppress the formation of larger-scale folded, misfolded and aggregated structures. The observations support a model in which Hsp90 alters client conformations directly by promoting local intra-chain interactions while suppressing distant ones. We conjecture that chain compaction may be central to how Hsp90 protects unstable kinases and receptor clients, regulates their activity, and how Hsp90 cooperates with Hsp70.

## Introduction

Heat shock protein 90 (Hsp90) is a chaperone that is highly conserved from bacteria to mammals and is essential in eukaryotic cells. It is involved in a wide array of cellular functions, ranging from protection against heat stress to signal transduction and protein trafficking (Taipale et al., 2010). Many of its functions involve cooperation with the Hsp70 chaperone system (Genest et al., 2011; Kirschke et al., 2014). Hsp90 interacts with many co-chaperones in eukaryotes (Echeverria et al., 2011; Mayer and Le Breton, 2015) whereas no co-chaperone has been identified for the *Escherichia coli* Hsp90, also termed HtpG (high temperature protein G). Two Hsp90 monomers form a V-shaped dimer that undergoes large conformational changes while progressing through the ATPase cycle (Fig. 1a). The Hsp90 dimer populates distinct conformations, ranging from a fully extended to a closed intertwined state, which can coexist in an equilibrium that depends on nucleotide, pH, and osmolyte conditions (Krukenberg et al., 2009; Shiau et al., 2006; Street et al., 2011). The closed intertwined state is poised for ATP hydrolysis.

**Fig 1.**
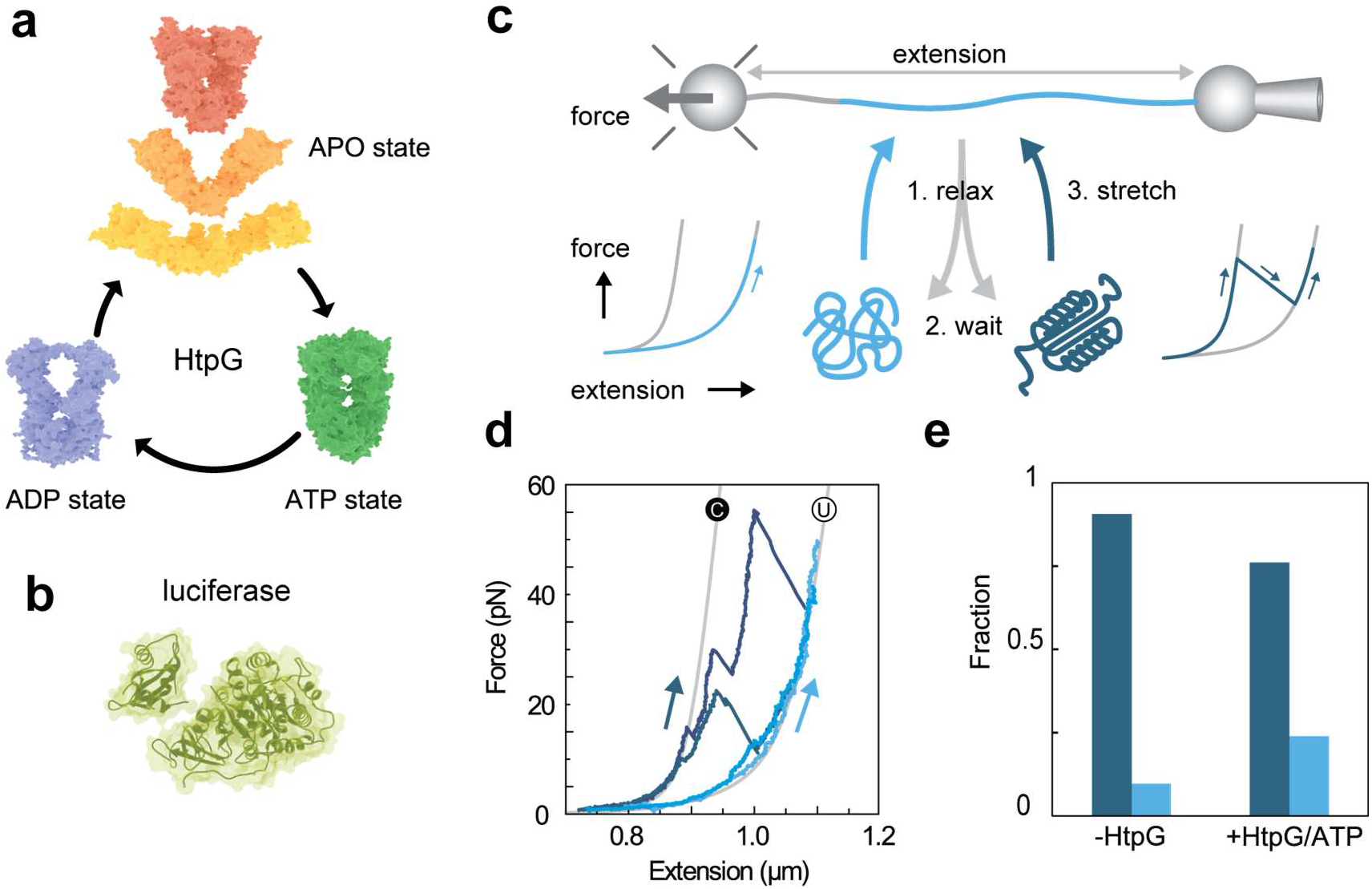
HtpG stabilizes the unfolded state of Luciferase. (a) Possible conformations assumed by bacterial Hsp90 (HtpG) when progressing through its ATPase cycle, ADP state 2IOP (Shiau et al., 2006), ATP state (*S. cerevisiae*) 2CG9 (Ali et al., 2006), APO states middle: 2IOQ (Shiau et al., 2006), APO states top and bottom (Krukenberg et al., 2009) (b) Firefly Luciferase structure 1LCY (Conti et al., 1996). (c) Schematic diagram of assay to measure stabilization of unfolded state. Luciferase is tethered between two beads using DNA linkers and fully unfolded and extended (top), relaxed to 0 pN for 5 sec. Subsequent stretching shows that the chain has remained unfolded (right, light blue) or adopted tertiary structure, as shown by an initially more compact state that unfolds in discrete steps (left, dark blue). Measured traces are shown in panel d, fractions of these two cases in panel e. (d) Stretching traces, taken from cycles as described in panel c, showing the chain remained unfolded (light blue) or adopted tertiary structure (dark blue). (e) Fraction of cycles showing protein chain remained unfolded (light blue) or adopted tertiary structure (dark blue). See panel c for description. Conditions are no HtpG (N=53 cycles), and 1 μM HtpG with 1 mM ATP (N = 126 cycles), see methods for details.

Hsp90 is thought to interact preferentially with unstable and intrinsically disordered protein regions (Schneider et al., 1996). For example, the intrinsically disordered protein Tau was found to bind an open conformation of Hsp90 in an ATP-independent manner, whereas ATP did affect Hsp90 conformation of pre-formed Tau-Hsp90 complexes (Karagoz et al., 2014). The intrinsically disordered ribosomal protein L2 accelerates ATP hydrolysis by Hsp90, suggesting that L2 binding can stimulate Hsp90 closure (Motojima-Miyazaki et al., 2010). Bacterial Hsp90 was found to bind a locally structured region of the partially folded model protein Δ131Δ (Street et al., 2014). Hsp90 was reported to bind the glucocorticoid receptor ligand-binding domain in open and closed conformations, though more efficiently in the former (Lorenz et al., 2014). In contrast to Δ131Δ and L2 however, binding here slowed down Hsp90 closure and decelerated ATP hydrolysis. Overall, these findings indicate that substrate binding affects and depends on Hsp90 conformation.

In contrast, it remains poorly understood how Hsp90 affects substrate conformations, which is central to understanding Hsp90 function. Bacterial Hsp90 can aid the Hsp70 system in refolding and reactivating misfolded luciferase (Genest et al., 2011; Moran Luengo et al., 2018). Eukaryotic Hsp90 can reactivate luciferase together with Hop/Sti1 (Johnson et al., 1998), and modulate the activity of steroid receptors in conjunction with the Hsp70 system and different co-chaperones (Morishima et al., 2000). Hsp90 can facilitate *de novo* folding in the cellular context, where various additional factors are present (Thomas and Baneyx, 2000). How protein chains are affected by Hsp90 alone is incompletely resolved (Street et al., 2014). Addressing this issue is technically challenging. The unstable and intrinsically disordered nature of Hsp90 substrates suggests that during their interaction with Hsp90, they may be characterized by conformational ensembles rather than distinct conformational states. Moreover, the diverse possible binding sites on the Hsp90 surface indicates the possibility of additional conformational heterogeneity and dynamics that are difficult to characterize.

Here we study this issue by manipulating individual luciferase molecules, as well as maltose binding protein and the glucocorticoid receptor, using optical tweezers. Our aim is to study how interactions with bacterial Hsp90 (HtpG) locally affect the conformation of unfolded substrates, rather than to investigate how Hsp90 assist in folding, which involves Hsp70 and other cofactors. Local structures that form in unfolded chains and hence change the distance between the N- and C-termini can be followed using optical tweezers for individual proteins at nanometer resolution, even if transient and heterogeneous. Our experiments showed that HtpG can produce a gradual compaction in unfolded luciferase substrates, as well as sudden step-wise contractions, in an ATP-stimulated manner, down to dimensions that approximate the fully compacted state. The HtpG interactions were also found to suppress entry into a kinetically trapped state and aggregation between proteins. The data indicate that HtpG promotes local intra-chain contacts within client protein chains while suppressing distant contacts, and hence induces a spectrum of transiently stable local conformations and overall chain compaction. This function may also have important implications for the interplay between Hsp90 and Hsp70 (Kirschke et al., 2014; Rodriguez et al., 2008; Sharma et al., 2010). More generally, unassisted spontaneous protein chain compaction or collapse is considered critical to protein states including phase separation (Dill, 1985; Kim and Baldwin, 1982). The promotion of compaction, which has thus far not been considered as a chaperone function, may hence be of general relevance to protein quality control.

## RESULTS

### Stabilization of the unfolded state by Hsp90

We surmised that one effect of Hsp90 could be the stabilization of unfolded substrate states, as the ability of Hsp90 to bind unfolded regions can compete with tertiary structure formation in the polypeptide chain. To investigate such stabilization of unfolded states by bacterial Hsp90 (HtpG) at the single-molecule level, we tethered a luciferase protein (Fig 1b) between two polystyrene beads via a DNA linker, and unfolded it fully by mechanical stretching (Fig. 1c). In these experiments, we continuously measure the distance between the beads, also referred to as the extension, as well as the force within the protein-DNA tether. Next, we relaxed the unfolded chain, waited at 0 pN for 5 s, and then stretched to assess whether it had remained unfolded or had formed tertiary structure (Mashaghi et al., 2014). For unfolded chains, stretching traces follow approximately a force-extension curve for a non-interacting polypeptide chain (of the length of luciferase) tethered to DNA, which is well-predicted by the worm-like chain (WLC) model (Fig. 1d, light blue traces) (Marko and Siggia, 1995). For chains with tertiary structure, the stretching data initially display compact conformations, which subsequently show discrete unfolding transitions (Fig. 1d, dark blue traces). In the absence of HtpG, we found that in 9% of the relaxation-stretching cycles, the chain remained in or near the unfolded state (Fig. 1e, light blue bar). In the presence of HtpG and ATP, the first stretching curves for newly tethered luciferase proteins were similar to the curves without HtpG, indicating a lack of interaction. After full unfolding, we similarly continued with cycles of relaxation, waiting at 0 pN for 5 s, and stretching. Now, a larger fraction (24%, p < 0.05, Fig. 1e, light blue bar) of the cycles showed that the chain remained in or near the unfolded state. These observations suggested that HtpG bound the unfolded chains, which is consistent with previous reports (Karagoz et al., 2014; Street et al., 2011), and could stabilize the unfolded state.

### Protein chain compaction by Hsp90

The relaxation data showed another type of effect on the unfolded luciferase substrates. Theory indicates that during relaxation to low force, an extended non-interacting protein chain coils up as described by the WLC force-extension curve (Fig. 2a, blue curve). Unfolded luciferase chains without HtpG present typically displayed that behaviour, while showing deviations at lower forces (Fig. 2b, below 5pN). In presence of HtpG and ATP, relaxation data deviated more strongly, with chains compacting when the force decreased below 25 pN, as indicated by extensions smaller than the unfolded-chain WLC model. We observed chains that became similarly compact as fully folded states for forces below a few pN, with force-extension data following the WLC model for the folded state (Fig. 2c-d). The compaction progressed gradually with decreasing force, while displaying small step-wise contractions down to the resolution limit of about 10 nm (Fig. 2d). Overall, these data indicated a gradual compaction of the protein chain.

**Fig 2.**
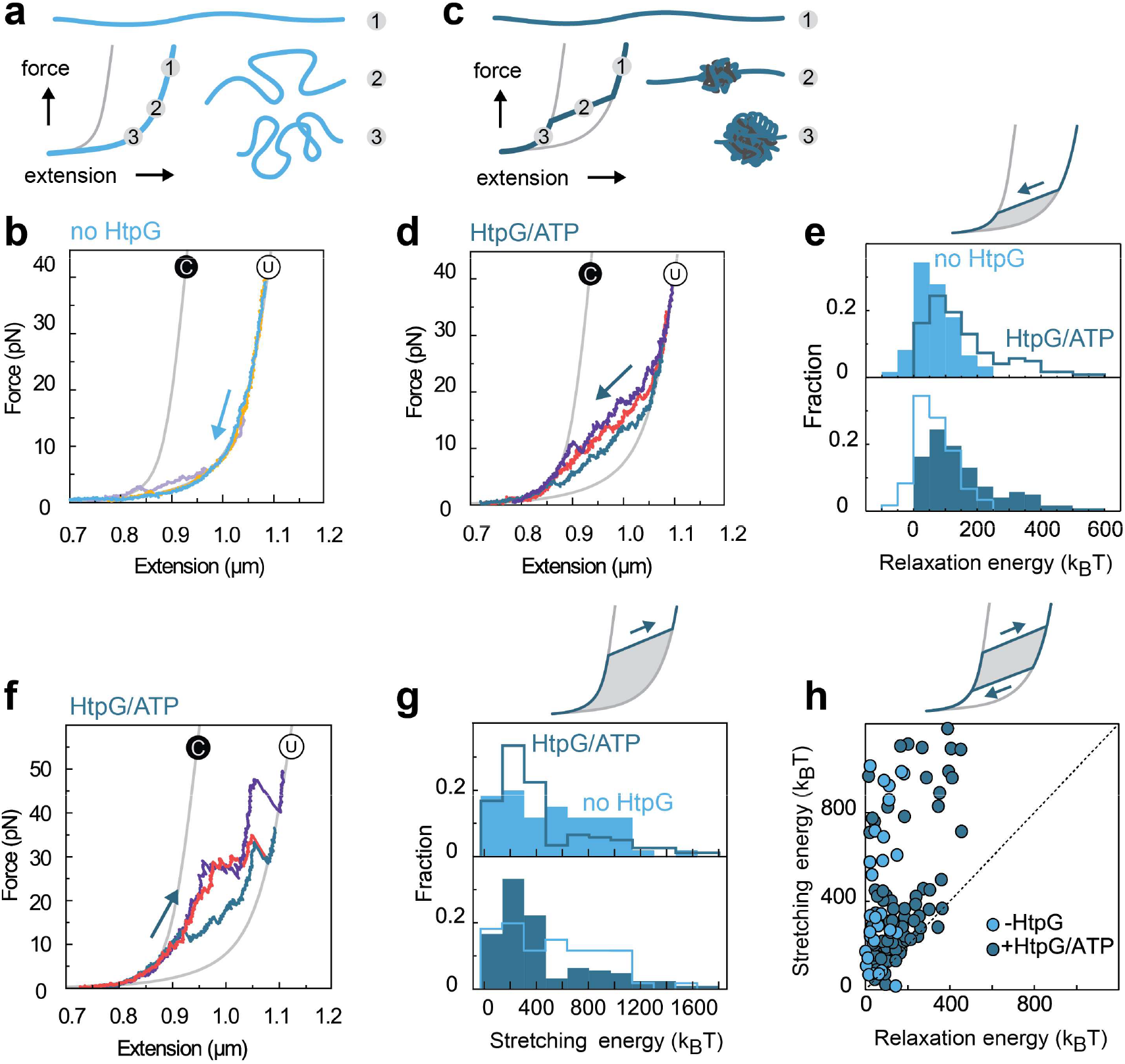
HtpG compacts unfolded Luciferase chains. (a) Cartoons of a non-interacting chain that coils up when the force is decreased from 1 to 3, and corresponding force-extension curve. (b) Measured force-extension data for unfolded luciferase that is relaxed from high to low force, from different relax-stretch cycles and different molecules. (c) Cartoons of a chain that collapses to a compact state when the force is decreased from 1 to 3, and corresponding force-extension curve. (d) Measured force-extension data for unfolded luciferase that is relaxed from high to low force in presence of HtpG and ATP, from different relax-stretch cycles and different molecules. (e) Histogram of observed relaxation energies in the absence (light blue, N = 63 traces) and presence of HtpG and ATP (dark blue, N = 125 traces), as quantified by the surface area in between the relaxation trace and the theoretical worm-like chain model (top). (f) Measured stretching data from low to high force in presence of HtpG and ATP, from different relax-stretch cycles and different molecules. (g) Histogram of stretching energy, in the absence (light blue, N = 63 traces) and presence of HtpG and ATP (dark blue, N = 125 traces) as quantified by the surface area in between the stretching trace and the theoretical worm-like chain model (top). (h) Hysteresis quantification. Relaxation energy against the stretching energy of the subsequent stretching trace. Distance from the diagonal quantifies the hysteresis of the cycle. Points on the diagonal reflect a lack of hysteresis. See also Figure S1.

To quantify the compaction effects, we determined the surface area under the relaxation traces (Fig. 2e). We are interested in deviations from a non-interacting chain, and hence we subtracted the area corresponding WLC behaviour, as well as that of the DNA linker (Methods). Note that this surface area reflects mechanical work. Here we refer to it as the ‘relaxation energy’, but note that it does not signify an equilibrium (free) energy. For the relaxation traces of luciferase without HtpG, this analysis showed a small but non-zero relaxation energy that extended up to 250 k_B_T (Fig. 2e, light blue). The order of magnitude of these effects are consistent with previous Atomic Force Microscopy pulling studies of the hydrophobic collapse of isolated protein chains (Walther et al., 2007). In line with the observed compaction, the distribution of the relaxation energy increased with HtpG and ATP now extended to 600 k_B_T (Fig. 2e, dark blue). Thus, HtpG induced contractions in the protein chain against a counter-acting force.

Next, we tested whether human Hsp90 (hHsp90) also induced chain compactions in one of its clients, the glucocorticoid receptor ligand binding domain (GRLBD). We found relaxation energies that were predominantly below 15 *k_B_T* in the absence of hHsp90 (Fig. S1). In the presence of hHsp90 and ATP, the relaxation energies were predominantly above 15 *k_B_T*, and the histogram now displayed a shoulder extending to up to 75 *k_B_T* (Fig. S1). Thus, hHsp90 promoted a contraction in the GR chain despite the counter-acting forces acting within it. Overall, these data indicated that both bacterial and hHsp90 can compact protein chains.

#### Hsp90 lowers the kinetic barrier to compacted states

In addition to the area under the relaxation trace which estimates the compaction energy, one may also quantify the area under the stretching traces (Fig. 2f). For luciferase, this stretching energy was found to decrease on average in the presence of HtpG (Fig. 2g, *p* < 0.05). Thus, whereas HtpG increased the relaxation energy, it decreased the stretching energy. These opposing trends indicated that the resulting compacted chain conformations, which are possibly complexed with HtpG, are less resistant to force than the folded conformations in absence of HtpG. The gradual compaction was also found to display small local steps, which likely involve few residues that are close together along the protein chain. In contrast, larger global folding transitions often involve many residues that may be far apart along the protein chain.

The notion of hysteresis can help to further discuss and explain this point. In general, hysteresis informs on the history-dependence of a system. A system is non-hysteretic when its current state does not depend on its previous state. For example, for a theoretical non-interacting chain undergoing relaxation-stretching cycles, the measured extension depends on the currently applied force as given by the WLC model, but not on previously applied forces. In such cases, the relaxation and stretching curves overlap, and the relaxation and stretching energies are identical. A different situation arises when considering (un)folding, for instance. As the tension is *increased* to a certain force, a chain that was folded may remain folded, but when the tension is *decreased* to that same force, the chain that was unfolded may remain unfolded. The delay in (un)folding is relevant here: both processes take time, which preserves previous states. The system is then not in thermodynamic equilibrium, and the difference in relaxation and stretching energies indicates the hysteresis. A limited hysteresis typically indicates low kinetic folding barriers and hence fast folding, as observed for small folds that exhibit limited cooperativity between residue-residue contacts (Jagannathan and Marqusee, 2013). Conversely, larger tertiary structures with complex folds and many cooperative contacts tend to exhibit large kinetic barriers, slow folding, and significant hysteresis (Shank et al., 2010).

Here we determine the hysteresis by the difference between the relaxation and stretching energies of each relax-stretch cycle, as assessed by plotting one against the other (Fig. 2h). The distance from the diagonal then quantifies the hysteresis. In the absence of HtpG, the cycles typically were separated from the diagonal by a distance of order 100 K_B_T, which indicated substantial hysteresis, and is expected for chain that folds (Fig. 2h, light blue points). In presence of HtpG and ATP, some cycles also displayed hysteresis, but many points were now close or on the diagonal, indicating limited to no hysteresis (Fig. 2h, dark blue points). The relaxation (and stretching) energies were up to 400 K_B_T in these non-hysteretic cycles. Thus, while the resulting states were compacted like folded states, they displayed a lack of hysteresis. Such reversibility has been observed in the formation and disruption of collapsed states in individual hydrophobic polymers (Li and Walker, 2011). Taken together, the decrease of hysteresis in presence of HtpG suggests that Hsp90 lowers the kinetic barrier to compacted states.

### Suppression of misfolding by HtpG

The data so far indicated that HtpG can promote the formation of compacted structures of dimensions down to similar order as fully folded states, within the measurement limits of our experiments. This compaction was gradual and displayed small steps (Fig. 2d). Consistent with the reversibility observed in the previous section, the stretching traces also displayed similar gradual and small step features (Fig. 2f), and were hence distinct from the stretching traces in the absence of HtpG (Fig. 1d, dark blue traces). These data are consistent with previous bulk data showing that HtpG alone does not refold luciferase, and rather requires cooperation with DnaK (Moran Luengo et al., 2018).

A related issue is whether Hsp90 affects entry into kinetically trapped misfolded states along the folding pathway. Luciferase is known to adopt non-native states by repeated freeze-thaw cycles (Genest et al., 2011; Johnson et al., 1998). Luciferase has also been shown to adopt kinetically trapped states in mechanical relaxation-stretching cycles (Mashaghi et al., 2014). During the 5 s waiting time at 0 pN in stretching and relaxation cycles, and in the absence of chaperone, luciferase indeed formed compact structures, termed X states, that are characterized by a protein length (the length of the unstructured part of the protein) of about 105 nm (13% of the cycles, Fig. 3a and 3b, red state). These states can be distinguished from larger (I1) and smaller (I2) intermediate structures (Fig. 3a and 3b). These X states have been identified as kinetically trapped misfolded states, as they have a significantly longer lifetime (minutes) and a maximally sustained force (43 pN on average) that is larger than other states of intermediate size (seconds and 24 pN respectively) (Mashaghi et al., 2014). This maximally sustained force equals the unfolding force when unfolding occurs, or the largest measured force when unfolding does not occur during stretching. In the presence of HtpG and ATP, the formation of misfolded X-state structures was almost abolished: their frequency was reduced more than 10-fold (from 13% to 0.8% of the cycles, *p* < 0.05, Fig. 3c). The other folded states including I1 and I2 did not show such a significant decrease (Fig. 3c). Overall, the data indicated that HtpG suppressed luciferase misfolding.

**Fig 3.**
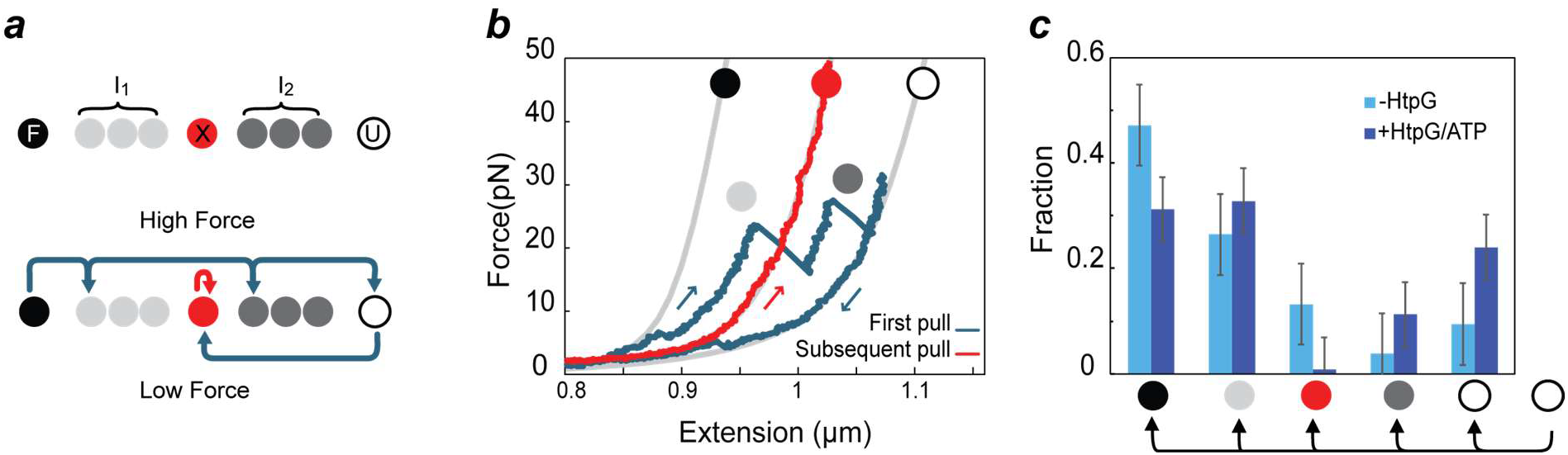
Misfold suppression by HtpG. (a) Schematic representation of observed luciferase states ordered by protein length, ranging from fully compact (F, black) to fully unfolded (U, white) state (top panel). Also indicated are a misfolded state (X, red), and states that are more compact (I_1_, light grey) or less compact (I_2_, dark grey). Bottom panel: transitions between states observed in panel b during stretching (top arrows) and in relaxed states (bottom arrows). (b) Stretching and relaxation (blue traces), waiting at 0 pN for 5s, and stretching (red trace). Data shows entry into state (X, red) of about 105 nm, which is stable over minutes and resists high forces, and corresponds to a misfolded state (Mashaghi et al., 2014). Grey lines are worm-like chain theoretical predictions of F and U states. (c) Corresponding fraction of observed entry into the different states indicated in panel a, in the absence (light blue, N = 53) and presence (dark blue, N = 126) bacterial Hsp90 (HtpG, 1μM). Cycles consist of relaxing chains in the fully unfolded state (right-most white circle), followed by a 5 s waiting time at 0 pN that provides an opportunity to adopt a different folded state. The latter is quantified in subsequent stretching, as illustrated in panel b. The data shows that HtpG suppresses entry into state X more strongly than the other states. Error bars represent the standard error calculated by assuming each population is independent. See also Figure S2.

To assess whether this misfold suppression by HtpG could occur beyond luciferase, we considered a different protein system. We focused on a construct composed of four maltose binding protein monomers arranged head-to-tail (4MBP), which has been used as a model system for protein misfolding and aggregation (Bechtluft et al., 2007; Mashaghi et al., 2013b). Stretching this 4MBP construct for the first time in the absence of HtpG results in an unfolding pattern that is characterized by four discrete events reflecting the unfolding of four MBP core structures (Fig. S2a and S2b). As shown previously (Bechtluft et al., 2007; Mashaghi et al., 2013b), after relaxation and waiting at 0 pN for 5 s, we observed compact structures that often could not be unfolded fully (Fig. S2c). The traces also showed steps larger than one MBP core, which thus indicated structures involving more than one MBP core (Fig. S2c and S2e). Together, these data indicated misfolding due to non-native contacts between different MBP cores. In the presence of HtpG and ATP, we found that the first stretching trace was similar to without HtpG, thus mirroring the lack of interaction in the first luciferase stretching traces. Subsequent stretching traces now were quite different. Tight misfolds that could not be unfolded were only rarely observed (Fig. S2d and S2e). Moreover, we now observed the discrete events reflecting the unfolding of precisely one MBP core (Fig. S2d). The presence of HtpG led to an about 3-fold increase of such events (Fig. S2e, from 4% to 13%, *p* < 0.05). Overall, these data suggested that HtpG suppressed global misfolding interactions between different cores while still allowing local interactions within single cores required for their refolding.

### Conformational changes promoted by HtpG are stimulated by ATP hydrolysis

HtpG undergoes conformational changes driven by an ATP hydrolysis cycle (Fig. 1a), which raises the question whether the observed effects depend upon it. In experiments with HtpG but without ATP (HtpG-APO), we found that the area under luciferase relaxation traces (Fig. 4a) was on average smaller than with HtpG and ATP (Fig. 4b, p < 0.05), indicating that HtpG-mediated compaction was stimulated by ATP presence. We found that for HtpG-APO the fraction of cycles where the chain remained in or near the fully unfolded state (31%, Fig. 4c) was about equal as for HtpG-ATP (24%), and hence higher than in the absence of HtpG (9%), consistent with HtpG-APO interacting with the Luciferase chain

**Fig 4.**
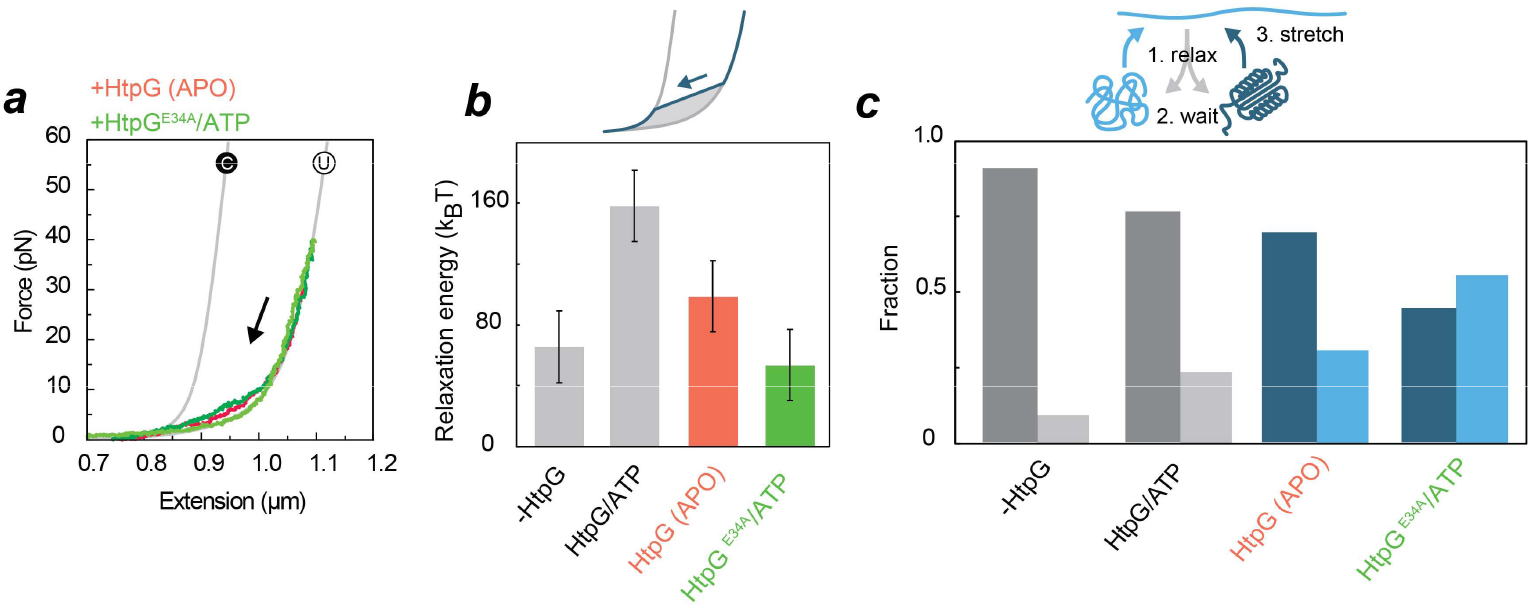
HtpG-promoted luciferase conformational changes are stimulated by ATP hydrolysis. (a) Relaxation traces with bacterial Hsp90 (HtpG, 1μM) and no ATP or HtpG-E34A with ATP. (b) Analysis of relaxation energy in the presence of HtpG/ATP (N = 125), HtpG (APO, N = 28), no chaperone (N = 63), and HtpG^E34A^/ATP (N = 18, Supplemental information). Error bars are SEM (c) Fraction of relaxation-stretching cycles that either did (dark blue) or did not (light blue) show the formation of tertiary structure (during relaxation and waiting at 0 pN for 5 s), with the formation of structure evidenced by discrete unfolding steps during subsequent stretching. Definitions as in Fig. 1e. See also Figure S3.

Next, we aimed to probe the role of ATP hydrolysis. ATP hydrolysis in Hsp90 proteins occurs within a conserved ATP binding site (Prodromou et al., 1997). In HtpG, a Glu to Ala mutation at residue 34 in this binding site decreases the rate of ATP hydrolysis about 10-fold (Genest et al., 2011; Graf et al., 2009). In the presence of HtpG^E34A^ and ATP, the stabilization of the unfolded state was most efficient, with 55% of cycles maintaining the chain in or near the fully unfolded state (Fig. 4c). These findings indicated that HtpG^E34A^ bound the unfolded luciferase. However, relaxation traces in the presence of HtpG^E34A^ displayed a low relaxation energy (Fig. 4a and 4b), and hence lacked the compaction that was seen for *WT* HtpG-ATP. These data indicated that HtpG^E34A^ with ATP interacted with the unfolded Luciferase chain but did not promote compaction.

We also observed a lack of compaction by *WT* HtpG and ATP in the presence of Radicicol (Fig. S3a,b), which is consistent with its ATPase inhibitory role. Overall, the data thus indicated that ATP hydrolysis stimulated HtpG-promoted compaction. We do note that for compaction of GRLBD by hHsp90 we did not find a stimulatory effect of ATP, but rather a small decrease in the relaxation energy from 42 K_B_T in absence of ATP (Fig. S1e) to 25 K_B_T in presence of ATP (Fig. S1a). Such differences in ATP dependence between HtpG and hHsp90 could be due to many factors, and may for instance be related to the stronger coupling of conformational change and the ATP cycle (Graf et al., 2014; Mickler et al., 2009; Ratzke et al., 2012), the client in question, or the absence of co-chaperones like Sti1/hop and p23 in the case of hHsp90.

## Discussion

Important advances have been made in revealing Hsp90 conformational changes throughout the ATP cycle and in response to interactions with its substrates, which are often structurally unstable or have an intrinsically disordered character (Ali et al., 2006; Graf et al., 2009; Krukenberg et al., 2008; Krukenberg et al., 2009; Mickler et al., 2009; Shiau et al., 2006; Street et al., 2010). Conversely, how the conformations of these unfolded substrate regions are affected by interactions with Hsp90 has remained largely unexplored. Work on the intrinsically disordered substrate Tau has shown it has a diverse and heterogeneous Hsp90-client binding interface (Karagoz et al., 2014). Bacterial Hsp90 has been shown to bind a region of Δ131Δ that is thought to be locally structured (Street et al., 2011). However, whether and how Hsp90 causes changes in its substrates has remained incompletely understood. Resolving this issue is crucial to elucidating the mechanism of Hsp90 action. Here we aimed to address it by measuring forces and length changes in individual protein chains in response to interactions with Hsp90, using optical tweezers.

Our data shows that bacterial Hsp90 (HtpG) can induce a global compaction of extended client chains driven by local contractions, down to overall dimensions that are similar to folded states within our detection limit. More specifically, we found that: 1) HtpG can trap a fraction of the substrates in states without significant stable tertiary structure, or low contact order (Fig. 1e), which suggests binding to extended segments or other small secondary or tertiary conformers of marginal stability. It also shows that such binding, which is consistent with diverse previous studies discussed above, can be stable enough to compete with tertiary structure formation and maintain substrates in such states of low contact order. 2) The local contractions yielded a gradual and global compaction of the protein chain, down to dimensions of the folded protein (Fig. 2d and 2e). This implies the presence of (effective) forces that drive the compaction, and hence can overcome opposing forces that drive chain expansion. The latter expansion forces may for instance originate from chain entropy or interactions with other chaperones, and can delay folding (Dill, 1985). 3) ATP hydrolysis suppression by the E34A mutation in HtpG abolished the stimulated compaction (Fig. 4a-4b), but led to efficient stabilization of low-contact-order states (Fig. 4c). Thus, ATP hydrolysis suppression limited one interaction mode (that caused compaction) while maintaining or promoting another (that caused stabilization of low contact order states). A similar mutation in yeast Hsp90 has shown that ATP hydrolysis was not absolutely essential for cellular viability, although the growth rate was reduced (Zierer et al., 2016). Differences in mutational effects between bacterial and yeast Hsp90 could be related to the tighter conformational control by ATP in the former (Mickler et al., 2009; Ratzke et al., 2012), or may reflect compensatory effects by other chaperones *in vivo*, or may indicate that eukaryotic Hsp90 needs cochaperones to perform similar tasks. 4) Hsp90 suppressed entry of unfolded chains into misfolded and aggregated states (Fig. 3c). These findings are consistent with increased aggregation in heat-shocked ΔHsp90 cells (Thomas and Baneyx, 2000), and the suppression of heat-activated aggregation by Hsp90 (Jakob et al., 1995), though the role of Hsp90 in *de novo* folding remains scarcely addressed.

The observed gradual compaction, including its observed reversibility and lack of hysteresis, indicates a spectrum of heterogeneous and transiently stable local conformations with low kinetic barriers. It is possible that binding to the open HtpG conformation and subsequent transition of HtpG to the closed intertwined conformation (Fig. 1a) promotes substrate compaction, while HtpG that is constitutively in the intertwined ATP bound state does not allow for a conformational change that drives substrate compaction. The substrate may also be in equilibrium between more compact and less compact local states, and that HtpG stabilizes the more compact states. Furthermore, one may speculate that the promotion of local compact conformations, which is consistent with previous indications of binding to local structures (Street et al., 2014), underlies the suppression of misfolding and aggregation observed here. By binding local structures, Hsp90 may shield these segments from interactions with other segments, which are distant and hence more likely to belong to different domains, and thus also produce non-native structures. Indeed, interactions between domains are considered a general factor contributing to misfolding (Han et al., 2007; Tian and Best, 2016). The stimulation of local interactions has been proposed as a model to suppress non-native interactions between domains by trigger factor (Mashaghi et al., 2013b; Singhal et al., 2015), but could apply to other chaperones as well. Differences between trigger factor and Hsp90 are also notable. The observed suppression of misfolding in luciferase and MBP suggests that HtpG binding leads to a generic destabilization, whether they are misfolded or native-like conformations, whereas trigger factor was shown to stabilize partially folded states against forced unfolding (Mashaghi et al., 2013b). This difference could, for instance, indicate that HtpG competes more than trigger factor for intra-chain interactions that stabilize tertiary structures. More generally, the compaction of protein chains that we observed is considered central to folding as it can bring chain segments together that form native interactions (Dill, 1985; Kim and Baldwin, 1982).

One may further speculate that the compaction by Hsp90 helps explain suggested transfer from Hsp70. Hsp90 could counteract the binding of Hsp70s to extended chain segments and the associated local unfolding. These processes are thought to underlie the activation and inactivation of a number of kinases and receptors in Eukaryotes, and hence play a regulatory role in key signal transduction pathways (Kirschke et al., 2014; Mashaghi et al., 2016; Rodriguez et al., 2008; Sharma et al., 2010). Such compaction could shield hydrophobic sites recognized by Hsp70s, thus preventing rebinding of Hsp70 to unfolded client segments (Moran Luengo et al., 2018). Hsp90 mediated compaction may also promote recently observed binding of Hsp70 to partially folded structures and resulting stabilization (Mashaghi et al., 2016), and thus assist in progression towards the native state.

In summary, the study provides a direct observation of how Hsp90 affects an unfolded protein chain. We find that it does not only binds chain segments, but can promote the formation of structures with low kinetic barriers, and an overall compaction of the protein chain. The findings have implications for understanding how Hsp90 affects the formation and stability of tertiary structures, and the cooperation between different chaperone systems. We speculate that Hsp90 may have a more direct role in altering the structure and function of its kinase and receptor clients, beyond binding disordered regions and hence preventing structure formation.

## AUTHOR CONTRIBUTIONS

A.M., F.M, E.K., M.P.M., G.K., and S.J.T. conceived and designed the research; G.K. designed and purified the MBP and luciferase protein constructs and the bacterial Hsp90; M.P.M. supplied the human Hsp90; E.K. designed and purified the GRLBD-DNA chimera; A.M., F.M., and E.K. performed the optical tweezers experiments; G.K., and M.P.M. supplied the chaperones; A.M., F.M, E.K., and S.J.T. analysed the data; and F.M, A.M., E.K., M.P.M., G.K., and S.J.T. wrote the paper. Work in the group of M.P.M was supported by the Deutsche Forschungsgemeinschaft (DFG) project ID 422001793 (MA1278/7-1)

## ACKNOWLEDGMENTS

Work in the group of S.J.T. is supported by the “Netherlands Organization for Scientific Research” (NWO). We thank D. P. Minde and Md. A. Kamal for fruitful discussions, and M. Avellaneda for help with preparing protein structure illustrations.

## EXPERIMENTAL PROCEDURES

### STAR Methods

#### Contact for Reagent and Resource Sharing

Further information and requests for resources and reagents should be directed to and will be fulfilled by the Lead Contact, Sander Tans.

#### Experimental Model and Subject Details

For the expression of avi-luci-4myc and 4MBP, E. coli BL21 (DE3) was used, for ybbR-luci-ybbR and GR-LBD T7 express (New England Biolabs), for HtpG E. coli MC4100 ΔhtpG∷kan, for Hsp90β E. coli BL21(DE3)Star/pCodonPlus (Invitrogen).

#### Method Details

##### Expression and purification of Luciferase

Luciferase was expressed and purified as follows (Mashaghi et al., 2014). Avi-luci-4myc was produced as hybrid protein consisting of an Ulp1-cleavable N-terminal His_10_-SUMO tag followed by an AviTag, luciferase and four consecutive myc-tags at the C-terminus. Overexpression of the Avi-luci-4myc-encoding gene was performed in *E. coli* BL21(DE3) cells harboring the BirA encoding plasmid pBirAcm (Avidity, LCC, Aurora, Colorado, USA) in LB medium supplemented with 20 mg/l Biotin, 20 mg/l Kanamycin, 10 mg/l Chloramphenicol, 0.1 mM IPTG at 20°C for about 20 hours. Cells from 1.5 l culture volume were lysed in buffer L containing 50 mM NaPO_4_ pH 8, 0.3 M NaCl, 10% glycerol, 2 mM β-mercaptoethanol. The lysate was cleared from cell debris by centrifugation at 35.000 g for 30 min and incubated for 1 hour with 2 g Ni-IDA matrix (Protino; Macherey-Nagel, Düren, Germany). The matrix was washed extensively with buffer L and bound protein was eluted in buffer L containing 250 mM imidazole. Eluate fractions containing the hybrid protein were pooled, His_6_-Ulp1 protease was added and dialyzed overnight at 4°C in buffer L. The next day, the protein mixture was subjected to a second Ni-IDA purification to remove the His-tagged protease and the His_10_-SUMO fragment and flow-through fractions containing purified Avi-luci-4myc were concentrated using Vivaspin concentration columns (Vivaproducts, Inc. Littleton, MA).

Complementary to the Avi-luci-4myc hybrid, an additional construct was produced in which Luciferase is flanked on both sides with the ybbR sequence (Yin et al., 2006). Overexpression was done in T7 express cells. The cultures were grown at 30°C to OD_600_ = 0.6 and expression was induced with 0.4 mM IPTG at 18°C for 20 h. Cells were lysed with a microfluidizer (EmulsiFlex-C3, Avestin) in lysis buffer L. The lysate was clarified by centrifugation (95000 g, 1h) and incubated with Ni-IDA matrix (Protino, Machery-Nagel) for 1 hour. After incubation, the matrix was washed with buffer L with 10 mM or 20 mM imidazole added and eluted with 250 mM imidazole

##### Expression and purification of HtpG

HtpG and HtpG^E34A^ were expressed as C-terminal His_10_-fusions in E. coli MC4100 ΔhtpG∷kan, and purified by Ni-IDA-chromatography (Protino, Macherey-Nagel) and anion-exchange chromatography (Resource™ Q, GE Healthcare). The purified proteins were checked to be nucleotide-free by anion-exchange chromatography (Resource™ Q) and by UV detection by 254 nm.

##### Expression and purification of human Hsp90β

Human Hsp90β was produced from the bacterial expression vector pCA528 (Andreasson et al., 2008) as fusion proteins with an N-terminal His6-Smt3 tag in the E. coli strain BL21(DE3)Star/pCodonPlus (Invitrogen). The cultures were grown to OD_600_ = 0.8-1.0 and expression was induced with 0.5 mM IPTG at 25°C overnight. Cells were lysed by a microfluidizer (EmulsiFlex-C5, Avestin) in lysis buffer A (40 mM HEPES/KOH pH 7.5, 100 mM KCl, 5 mM MgCl2, 10% glycerol, 4 mM β-mercaptoethanol) and 5 mM PMSF, 1 mM Pepstatin A, 1 mM Leupeptin and 1 mM Aprotinin. The lysate first was clarified by centrifugation (30,000 g for 30 min), and then incubated with Ni-IDA matrix (Protino, Macherey-Nagel) for 30 min. After incubation, the matrix was washed with buffer A and bound protein eluted with buffer A containing 250 mM imidazole. The eluted fusion proteins were supplemented with Ulp1 protease, which cleaved the His_6_-Smt3 tag, and the mixture was dialyzed overnight against buffer A containing 20 mM KCl. Cleaved recombinant proteins were recovered in the flow-through fractions after a second incubation with Ni-IDA matrix whereas the N-terminal His6-Smt3 tag and Ulp1 remained on the column. Proteins were further purified by anion-exchange chromatography (Resource™ Q; GE Healthcare) with a linear gradient of 0.02–1 M KCl, fractions of eluted proteins were subjected to Superdex 200 16/60 Prep grade column in storage buffer (40 mM HEPES/KOH, pH 7.5, 50 mM KCl, 5 mM MgCl2, 10% glycerol, 4 mM β-mercaptoethanol). The purity and molecular mass were verified by SDS–PAGE and HPLC-electrospray mass spectrometry, confirming the correct primary sequence containing only the N-terminal start-methionine.

##### Expression and purification of 4MBP

N-terminally biotinylated 4MBP C-terminally fused with 4 Myc tag sequences were produced in *E. coli* BL21(DE3) as hybrid proteins consisting of an N-terminal Ulp1-cleavable N-terminal His_10_-SUMO sequence followed by an AviTag sequence (Avidity, LCC, Aurora, Colorado, USA), facilitating in vivo biotinylation and four consecutive C-terminal Myc-tag sequences. Proteins were purified from *E. coli* BL21(DE3) cells harboring pBirAcm encoding the biotin ligase (Avidity, LCC, Aurora, Colorado, USA). For over-expression cells over-night cultures were diluted 1:100 in fresh LB medium supplemented with 20 mg/l Biotin, 20 mg/l Kanamycin, 10 mg/l Chloramphenicol, 0.2% glucose and incubated under vigorous shaking at 30°C. Expression was induced at OD_600_= 0.6 by addition of 1 mM IPTG for 3 h. Cells were chilled, harvested by centrifugation, flash-frozen in liquid nitrogen and stored at −70°C. Cell pellets were resuspended in ice-cold buffer A (20 mM Tris-HCl pH 7.5, 0.2 M NaCl, 1% Triton X-100, 1 mM PMSF) and lysed using a French Pressure Cell. The lysate was cleared from cell debris by centrifugation at 35.000 g for 30 min and incubated with Ni-IDA matrix (Protino; Macherey-Nagel, Düren, Germany) for 30 min at 4°C. The matrix was washed extensively with buffer A and bound hybrid proteins were eluted in buffer A containing 250 mM imidazole. The eluate was supplemented with His_6_-Ulp1 protease and dialyzed overnight at 4°C in buffer D (20 mM Tris-HCl pH 7.5, 0.2 M NaCl). Following dialysis coupled with Ulp1 digestion, His_6_−Ulp1 protease and the His_10_−SUMO fragment were removed by incubation with Ni-IDA matrix. The unbound MBP or 4 MBP that remained in the unbound fraction was then loaded on Amylose resin (New England Biolabs) previously equilibrated in buffer D, washed with cold buffer D and bound proteins were eluted in buffer D supplemented with 20 mM maltose. Elution fractions were dialyzed three times for 2 hours at 4°C in 100-fold excess volume of buffer S (20 mM Tris-HCl pH 7.5, 0.2 M NaCl, 1 mM EDTA). 4 MBP purifications in addition were subjected to size-exclusion chromatography using a Superdex 200 HiLoad 16/60 prep grade column. Purified proteins were concentrated using Vivaspin centrifugal concentrators, aliquoted, flash frozen in liquid nitrogen and stored at −70 °C.

##### Expression and purification of GRLBD

Human GRLBD-F602S (520-778), flanked with an AviTag (Beckett et al., 2008) on the N-terminal side and a Sortag (Popp et al., 2007) on the C-terminus, was produced in *E. coli* T7 express (New England Biolabs) as an MBP fusion in the pMAL-c2E vector in the presence of 100 μg/L Ampicillin, 17 μg/L Chloramphenicol, 20 mg/L D-biotin and 50 μM dexamethasone (Sigma-Aldrich). The protein was in-vivo biotinylated using BirA produced from the pBirAcm plasmid (Avidity LCC, Aurora, CO, USA). Expression of the GRLBD-encoding gene was induced with 0.5 mM IPTG at 15°C for 16 hours. The cells where harvested by centrifugation 4°C, 5000 g, dissolved in lysis buffer (wash buffer without ATP and with complete protease inhibitor cocktail (Roche) 20 min, and lysed using Emulsiflex C3 homogenizer (Avestin, Ottawa, Canada). Insoluble material was removed by ultracentrifugation at 100,000 g for 1h at 4°C. The lysate was affinity purified using amylose resin (New England Biolabs, Ipswich, MA, USA). Wash buffer was 50 mM Tris pH 8.3, 300 mM KCl, 5 mM MgCl_2_, 0.04% CHAPS, 1 mM EDTA, 10% glycerol, 50 μM dexamethasone, 2 mM ATP and 5 mM β-mercaptoethanol. GR-LBD was eluted with wash buffer supplemented with 10 mM D-Maltose. Using a PD-10 desalting column (GE healthcare) buffer was exchanged to storage buffer (50 mM Tris pH 8.3, 100 mM KCl, 5 mM MgCl_2_, 10% glycerol, 50 μM dexamethasone and 5 mM β-mercaptoethanol).

##### Coupling of oligos to GRLBD

5’NH2-labeled CAGGGCTCTCTAGATTGACT (IDT-DNA, Leuven, Belgium) was coupled with sulfo-SMCC (Thermo-Fisher) according to the manufacturer’s instructions resulting in an maleimide-labeled oligo. The product was purified by ethanol precipitation. A GlyGlyGlyCys peptide (Proteogenix, Schiltigheim, France), was dissolved in MeOH to 10 mM and added in a 6:1 ratio to 300 μM oligo-maleimide and incubated for 1h at 37°C. To drive the reaction to completion, 0.5 mM TCEP was added and the reaction mixture was incubated for 30 min at 37 °C. The reaction was stopped by addition of 0.1 M sodium molybdate (Kalia and Raines, 2007). The product purified by 2 ethanol-precipitation steps. The glycine-modified oligo is coupled to the GR using the Sortase A reaction (Antos et al., 2017). 7M Sortase A was expressed in *E. coli* from pet30b-7M SrtA (Addgene plasmid # 51141 was a gift from Hidde Ploegh) and purified according to Popp et al. (Antos et al., 2017). In order to remove the non-reacted glycine-modified oligo, a second affinity (amylose) purification was performed.

##### Coupling of oligos to Luciferase

The oligo, 5’-labeled with Coenzyme A (Biomers GmbH), SFP synthase (New England Biolabs) and purified ybbR-Luciferase-ybbR were incubated over night at 4°C. A second His-tag purification was done to remove the non-reacted Coenzyme A-modified oligo.

##### Coupling of DNA tether to GR-oligo and Luciferase-oligo

A 2.5 kbp DNA fragment was PCR amplified from the pUC19 plasmid (New England Biolabs) with a double digoxigenin-labeled primer (Biomers GmbH) on one side and a phosphoprimer on the other side and purified using the QIAquick PCR purification kit (Qiagen, Hilden, Germany). The phosphorylated strand is digested by Lambda exonuclease (New England Biolabs) for 2 hours at 37°C and purified using an Amicon 30 kDa MWCO filter (Merck, Darmstadt, Germany). The Deep Vent exo- DNA polymerase (New England Biolabs) and a 20 nt more upstream primer than the phosphoprimer from the PCR is used for the fill up of the second DNA strand creating a 20 nt overhang. This overhang is complementary to the 20 nt oligonucleotide sequence ligated to the C-terminus of GRLBD. The overhang DNA is added to the GR-oligo together with T4 ligase (New England Biolabs) and incubated for 30 min at 16°C followed by 30 min on ice. The resulting GR-DNA hybrid was flash frozen and stored at −80°C until measurement.

##### Coupling of DNA tether to Luciferase-oligo

A similar procedure was used for attaching two long DNA handles two ybbR-Luciferase-ybbR as for the GR-oligo. For this reaction, half of the DNA was a biotin labeled and the other half contained the (previously described) digoxigenin-labeled DNA. The biotin labeled DNA handles were made in the same way was as digoxigenin-labed DNA, but instead of a digoxigenin primer, a triple-labeled biotin primer (Biomers GmbH) was used.

##### Bead preparation Luciferase and 4MBP

We used commercially provided AntiDig-coated beads (Spherotech, 2μm diameter), Neutravidin-coated bead (Spherotec, 2.1 μm diameter) and prepared the AntiMyc-coated beads in-house. Anti-c-Myc (Roche) molecules were covalently linked to the carboxylated polystyrene beads (Spherotech) via Carbodiimide reaction (PolyLink Protein Coupling Kit, Polysciences Inc.). Briefly, 5 μl of 5% (w/v) 2 μm diameter carboxylated polystyrene microspheres were washed twice by pelleting at 13200 rpm (for 10 min) in a microcentrifuge tube and resuspending in coupling buffer (400 μl in first wash and 170 μl in second washing) (PolyLink Protein Coupling Kit, Polysciences Inc.). Then 20 μl of the freshly prepared EDCA solution (20 mg/ml; prepared by dissolving 1 mg EDCA in 50 μl coupling buffer) was added to the microparticle suspension and mixed gently end-over-end. After that, 20 μg of Anti-c-Myc was added and the mixture was incubated for 1 hr at room temperature with gentle mixing. Then the mixture was washed twice in 400 μl storage buffer. AntiMyc-coated beads were stored in 400 μl storage buffer (containing 0.05% BSA) at 4°C until use.

DNA-coated microspheres were made by mixing ~70 ng of dsDNA molecules and 1 μl AntiDig-coated beads in 10 μl HKM (50 mM HEPES, pH 7.6, 100 mM KCl, 5 mM MgCl_2_) buffer. After 30 minutes incubation on a rotary mixer (4°C), 1 μg of Neutravidin was added to the beads solution. Beads were again incubated for 15 minutes on a rotary mixer (4°C). The unreacted Neutavidin molecules were separated by pelleting the beads at 13200 rpm (for 10 min) in a microcentrifuge tube. After taking the supernatant, the beads were resuspended in 400 μl HKM buffer for use in optical tweezers experiments. Protein-coated microspheres were made by mixing ~10 μg of Luciferase molecules and 1 μl AntiMyc-coated beads in 10 μl HKM (50 mM HEPES, pH 7.6, 100 mM KCl, 5 mM MgCl_2_) buffer. After 30 minutes incubation on a rotary mixer (4°C), the beads were diluted in 400 μl HKM buffer for use in optical tweezers experiments.

For the ybbR-Luciferase-ybbR construct, the same procedure was used as for GRLBD, except that the measurements were done in HKM buffer.

##### Bead preparation GRLBD

The GR-DNA chimeras were tethered to the surface of 2.1 μm AntiDig-coated polystyrene beads in HMKDM buffer (50 mM HEPES pH 7.5, 5 mM MgCl_2_ and 100 mM KCl, 50 uM dexamethasone, 5 mM β-mercaptoethanol). After 40 minutes incubation on a rotary mixer (4°C), the beads were diluted in 175 μl HMKDM buffer for use in optical tweezers experiments supplemented with an enzymatic oxygen scavenger system (Swoboda et al., 2012). For experiments in the presence of chaperone, the HMKDM supplemented with Hsp90-β (2 μM), ATP (1 mM), phosphoenolpyruvate (3 mM), and pyruvate kinase (20 μg/mL).

##### Optical tweezers buffer conditions

The MBP and Luciferase experiments were carried out in HMK (50 mM HEPES, pH 7.6, 100 mM KCl, 5 mM MgCl_2_) buffer, supplemented with HtpG (1 μM), HtpG^E34A^ (5 μM), and ATP (1 mM). The GR experiments were carried out in HMKDM buffer (50 mM HEPES pH 7.5, 5 mM MgCl_2_ and 100 mM KCl, 50 uM dexamethasone, 5 mM β-mercaptoethanol), supplemented with Hsp90-β (2 μM), ATP (1 mM), phosphoenolpyruvate (3 mM), and pyruvate kinase (20 μg/mL).

##### Optical tweezer setup

Two optical tweezers set-ups were used. The first is a custom-built single trap optical tweezers (Mashaghi et al., 2013a; Moayed et al., 2013). Detection of forces on the trapped bead was performed using back focal plane interferometry. Forces were recorded at 50 Hz. A piezo-nanopositioning stage (Physik Instrumente) was used to move the sample cell and micropipette at a speed of 50 nm s^−1^. The beads were trapped in a flow chamber with three input channels: one containing AntiMyc-coated beads with the protein construct; one containing AntiDig-coated beads with the DNA linker and a central buffer channel in which the measurements were conducted. The second is a custom-built dual-trap optical tweezers. A single solid-state laser (IPG Photonics, 5 W) was split by polarization into two independently controlled traps. Forces were monitored in both traps separately using back-focal plane interferometry (Gittes and Schmidt, 1998) and bead displacement was calculated according to Δ*F* = −*κ*Δ*y*. The stiffness of the traps was determined by thermal calibration using the power spectrum method (Nørrelykke and Flyvbjerg, 2010). Data was acquired at 195 Hz.

#### Quantification and Statistical Analysis

##### Protein length determination

To determine measured protein lengths, we fitted measured force-extension data to a worm-like chain model (Wang et al., 1997) of a DNA-Polypeptide construct, with the polypeptide-contour length (*L_p_*) as the fitting parameter (S1). D and p indexes indicate DNA and polypeptide, respectively.

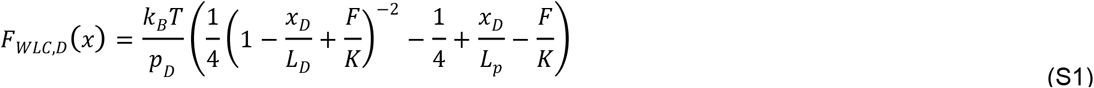

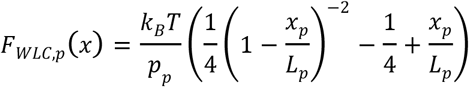

Here, *F* is the force, *x* is the extension (end-to-end distance), *L* is the contour length, *p* is the persistence length, *K* is the stretch modulus, k_B_ is the Boltzmann constant, and *T* is absolute temperature. The force-extension data was fitted considering the DNA linker and the polypeptide as two springs in series, i.e. *F* = *F*_*WLC,D*_ = *F*_*WLC,p*_; *x*_*Total*_ = *x*_*D*_+*x*_*p*_. We note the protein length is assumed to be equal to the contour length of the unfolded part of the protein chain, given the comparatively small contribution of folded structures to the measured length. Worm-like-chain parameters used in this study are the following: For the MBP and Luciferase-Myc experiments, K_D_=1200 pN, p_D_=45 nm, p_P_=1nm, L_P_ (luciferase)=198nm, L_P_ (4MBP)=480nm, L_D_=920nm was used. For the luciferase-ybbR experiments: p_D_ = 20 nm, K_D_ = 1250 pN, p_P_= 0.75 nm, L_P_ =187 nm, L_D_=906 nm. For the GRLBD measurements: p_D_ = 20 nm, K_D_ = 1200 pN, p_P_= 0.75 nm, L_P_ =95 nm, L_D_=906 nm was used.

##### Analysis of force-extension data

Before taking data on a particular tether, we performed several controls to confirm only a single tether is present. We check that the total unfolding of the proper expected length, and that the force-extension data is consistent with the WLC model (at higher forces). We also monitor that the tethers overstretch at 65 pN. At the end of experimentation on a tether, we check that it breaks in one clean break. For the analysis of Fig. 1e, 3c, and 4c, we followed previous work on the folding and unfolding of Luciferase alone (Mashaghi et al., 2014), which showed that the smallest partially refolded state of Luciferase unfolded with a contour length change of 31 nm. We divided the relax-stretch cycles into two categories: stretching curves that showed contour-length changes of less than 31 nm in total above 5pN (and hence have remained predominantly unfolded, light blue (or gray) bars Fig. 1e and 4c), and those showing contour length changes of more than 31 nm above 5 pN (dark blue (or gray) bars Fig. 1e and 4c). We observed chains that for a number of cycles did not refold, and then again refolded. This longer-term memory could be due to slow proline isomerization or residue rotational angle relaxation. There is the added possibility that HtpG remains associated, which can also lead to memory. The relaxation energy (Fig. 2e, 4b and S1a,c, and e) refers to the surface area in between an experimental force-extension curve during relaxation, and the theoretical WLC force-extension curve for the DNA tether connected to a polypeptide of appropriate length (equation S1) (Mossa et al., 2009). Similarly, we refer to the stretching energy (Fig. 2g) as the surface area in between an experimental force-extension curve during stretching, and the theoretical WLC force-extension curve for the DNA tether connected to a polypeptide of appropriate length.

## Notes

### Competing Interest Statement

The authors have declared no competing interest.

