## Supplemental Information for "Direct observation of Hsp90-induced compaction in a protein chain"

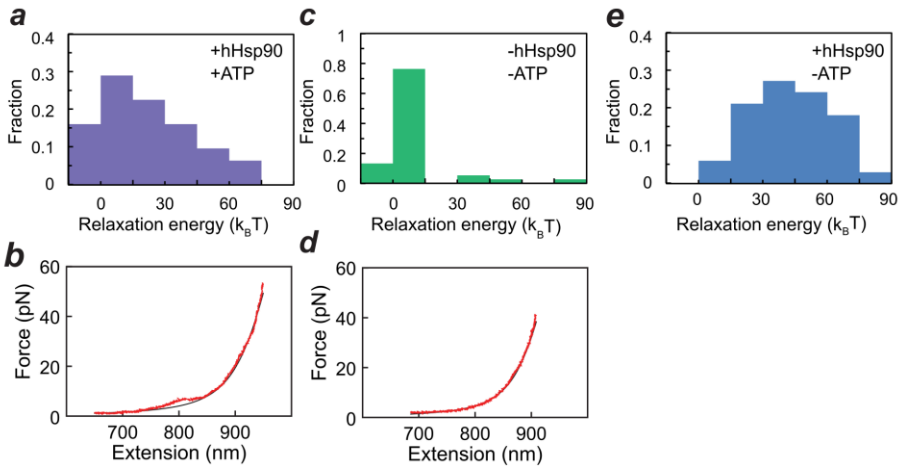


**Figure S1. Relaxation energy of GRLBD, related to Figure 2**. **(**a) Relaxation energy for GRLBD in the presence of 2 μM human Hsp90β (hHsp90) and ATP, as determined by the deviation from the worm-like chain behavior with an average value of 25±4 $k_BT$ (N = 27). (b) Corresponding relaxation curve (red) and worm-like chain model for non-interacting chains (black) with hHsp90 and ATP (see panel a for histogram). (c) Relaxation energy for GRLBD without hHsp90, as determined by the deviation from the worm-like chain behavior with an average value of 8±2 $k_BT$ (N = 38). (d) Corresponding relaxation curve (red) and worm-like chain model for non-interacting chains (black) without hHsp90 (see panel c for histogram). (e) Relaxation energy for GRLBD in the presence of 2 μM human Hsp90β (hHsp90) and absence of ATP, as determined by the deviation from the worm-like chain behavior with an average value of 42±3 $k_BT$ (N = 33)

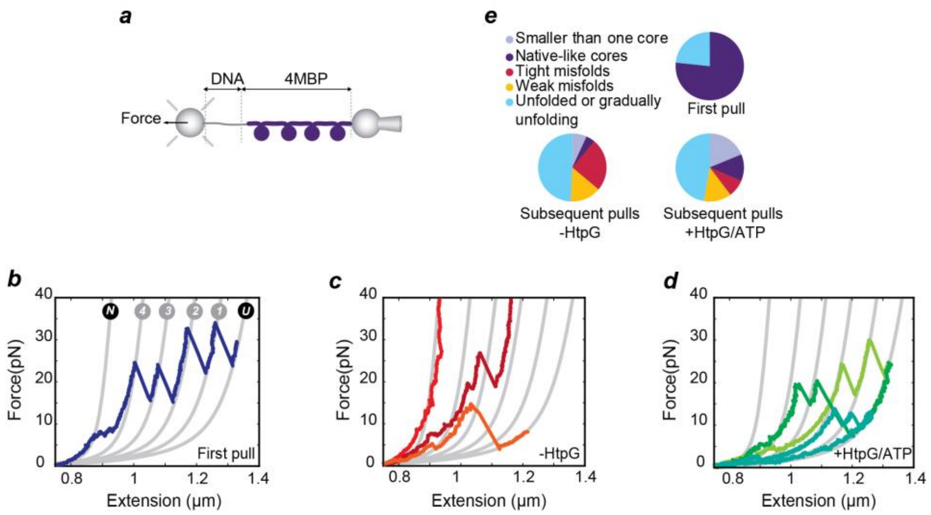


**Figure S2. HtpG Suppresses inter-domain misfolding, related to Figure 3.** (a) Schematic representation of a 4MBP molecule tethered between two anti-body-coated beads via a DNA linker. (b) Stretching curves for the 4MB construct, showing the unfolding pattern of natively-folded 4MBP. Gray lines represent the theoretical WLC characterizing the DNA-protein construct from fully-folded (N) to fully unfolded (U) state. After C-terminal unfolding (N→4), four native-like core unfolding events (4 →3 →2 → 1 → U) are observed. These first stretching curves on newly tethered proteins are similar in the presence and absence of HtpG, indicating a lack of interaction between HtpG and the native state (N=10). (c) Stretching curve after full unfolding and relaxation in the absence of HtpG. These subsequent stretching curves typically show either what we refer to as tight misfolds (that fail to unfold for forces up-to our maximum of 65 pN when the DNA melts) or weak misfolds (which do unfold but release chain segments exceeding one core structure, i.e. 92 nm, and hence must represent a structure involving more than one MBP repeat) (N=37). (d) Subsequent stretching curves after relaxation of fully unfolded proteins, in the presence of HtpG/ATP. The curves show that tight misfolds are suppressed while native-like folds are promoted (see last two steps in the light-green curves that match the lengths of two core domains unfolding) (N=23). (e) Corresponding fractions of different types of events.

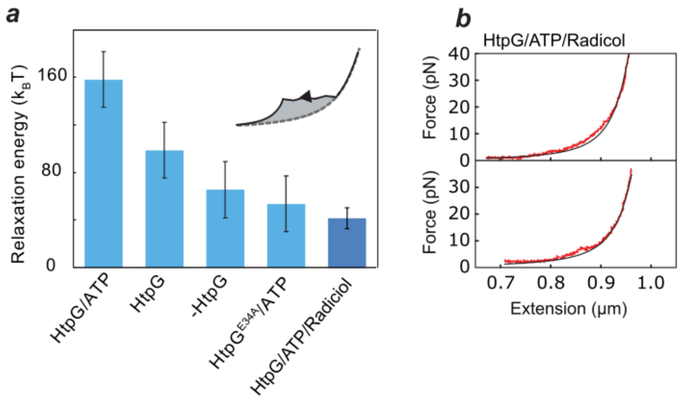


**Figure S3. Dependence on ATP hydrolysis and radicicol, related to Figure 4.** (a) Additional experiments to fig 4c, performed with 1 μM HtpG, 1 mM ATP and 1 μM radicicol. Light blue: relaxation energies from figure 4c as a reference. Dark blue: relaxation energy for HtpG/ATP/Radicicol (N = 31). Error bars are SEM (b) Examples of relaxation curves of luciferase with HtpG/ATP/Radicicol and worm-like-chain curve (black), used to calculate the relaxation energy in panel a. See Fig. 2d for relaxation traces in absence of Radicicol and presence of HtpG and ATP.
